# Stable Actin As Synaptic Tag

**DOI:** 10.1101/2025.02.10.637360

**Authors:** Mitha Thomas, Cristian-Alexandru Bogaciu, Silvio O. Rizzoli, Michael Fauth

## Abstract

According to the tagging and capture hypothesis, long-lasting long-term potentiation (LTP) requires protein synthesis and a synaptic tag, which is a synapse specific memory of the synapse with a so far unclear molecular or biophysical identity. Here we use an interdisciplinary approach to explore the hypothesis that interaction between the dynamics of actin and the spine geometry can provide such a memory. Using a mathematical model, we demonstrate that this implementation of the tag requires an increase in the stable, cross-linked pool of actin filaments, and is not possible without this stable pool. Using FRAP experiments, we show that such an increase in stable actin can indeed be observed after cLTP in vitro. Thus, the interaction between actin dynamics and spine geometry could indeed serve as a synaptic tag for LTP.

## Introduction

Long-term potentiation (LTP, [1]) of excitatory synapses changes the connectivity of neuronal networks and their ability to process information. Accordingly, LTP has been associated to learning and long-term memory formation in neuronal networks [2–4].

However, long-term potentiation can be expressed in different forms[5, 6]: as early LTP (E-LTP), which decays within hours, and as late LTP (L-LTP), which persists much longer. The most common model for the emergence of L-LTP – the synaptic tagging-and-capture hypothesis[5, 7] – states that late LTP requires two components: (1) a transient memory that the synapse experienced a plasticity inducing event, the so-called synaptic tag, and (2) *de novo* synthesis of plasticity-related proteins (PRPs), which are then translocated to the tagged synapse and give rise to L-LTP. If PRPs are missing, only E-LTP is observed. The molecular or biophysical identity of the synaptic tag is not completely clear[7, 8], but one molecule that is intimately related to the synaptic tag is the scaffolding protein actin[9]. Actin forms dynamic filaments that continuously polymerize at their barbed end and depolymerize at their pointed end (treadmilling). Additionally, various actin binding proteins (ABPs) organize the filaments into branched networks (ARP2/3 complex), sever the filaments (ADF/cofilin), or cap the barbed ends and prevent polymerization (capping protein)[10]. Moreover, the filaments can be distinguished into (at least) two dynamically distinct pools of actin: a fast-treadmilling, dynamic pool and a stable pool, in which filaments are bound to cross-linkers like *α-*actinin, drebrin, cortactin[11] or CaMKII[12], which slow down the filament dynamics [13]. These pools undergo massive reorganization during LTP, as the concentration of many actin-binding proteins and cross-linkers changes in a time-dependent manner [14, 15]. Yet, such transient changes in actin alone cannot account for the tag [7, 16, 17].

However, as actin forms the scaffold of the spine, its time-dependent reorganization also gives rise to massive changes in the geometry of the dendritic spines on which the excitatory synapses reside – most prominently an enlargement of the spine volume [18]. Thus, there is a complex interaction between actin pools and spine geometry, which may retain the information of a plasticity event longer than actin dynamics alone and would also be consistent with the tag being a “temporary structural state of the synapse”[7].

To understand this complex interaction, one can use mathematical modeling, which is based on, and informs, experimental measurements. Along this line, a variety of models describing the interaction between actin dynamics and spine geometry have been published[19–24] Yet, these models usually only consider the dynamic pool of actin filaments that drives the expansion of dendritic spines, and do not explicitly consider the existence or re-organization of the stable pool.

Accordingly, these models can only account for the first few minutes after LTP, where the initial enlargement of the spine takes place.

It is, however, unclear whether these models exhibit a persistent change of spine geometry and actin dynamic on the timescale of the synaptic tag, which is typically around one to two hours [5]. Moreover, while there is evidence that cross-linked filaments play a major role during [13] and after LTP[25], it is unclear whether the stable pool exhibits any alterations at this timescale.

Therefore, in this study, we follow an interdisciplinary approach combining theory and experiments to better understand the role of the stable actin pool in LTP. First, we test whether an experimentally constrained model[22] without a stable pool exhibits altered actin dynamics or spine geometry at the timescale of the synaptic tag, and find that this is not the case. We then experimentally evaluate the fraction of stable actin 30 minutes after cLTP, and find that it is increased by a factor of 2-3. Therefore, we adapt the theoretical model to include a dynamical stable pool and demonstrate that this refined model indeed exhibits altered spine geometry and actin dynamics on the timescale of hours, supporting the hypothesis that actin and spine geometry serve as a biophysical implementation of the synaptic tag.

## Results

### Model of dynamic actin and its modulation during LTP

We first test whether a model without a stable pool exhibits long-lasting perturbations of spine volume or actin dynamics after LTP. To this end, we exposed an existing model[21], which is known to match the dynamic of actual spines[22], to the characteristic changes in the concentrations of ABPs observed during LTP[14].

In the model, the spine membrane is described by a triangular mesh. Each point of the mesh, aside from points belonging to the PSD and the spine neck, moves according to the local balance between forces exerted by the membrane and by the actin network. The membrane force is derived from the Canham-Helfrich free energy formalism and accounts for the membrane resisting against changes in spine volume, surface area and surface curvature (Fig. 1A right, see Methods: Membrane force).

**Fig. 1.**
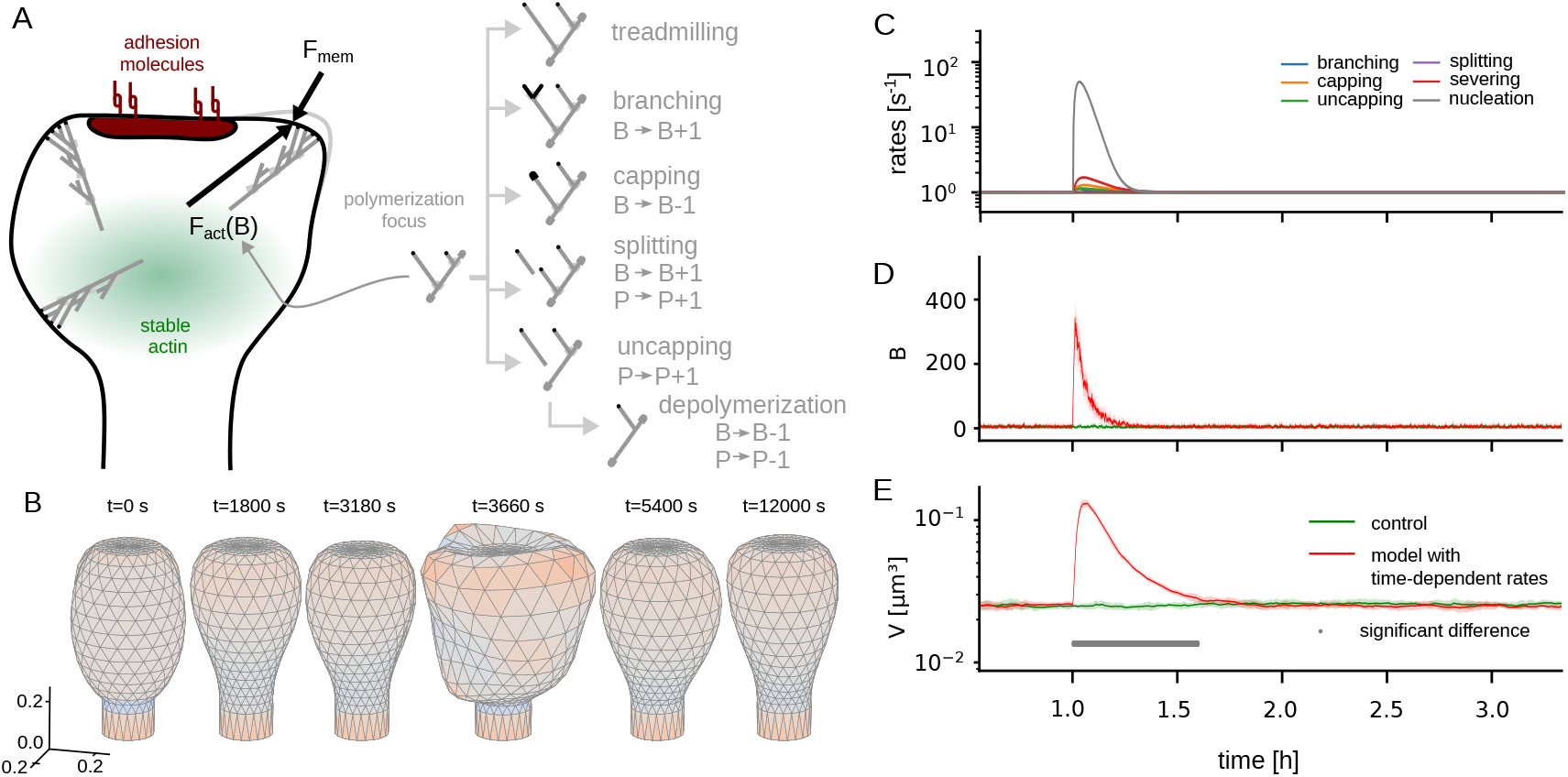
LTP-induced changes in actin dynamics and spine volume decay quickly. (A) Scheme of model components (B) Example spine shapes from simulation. Colors indicate the curvature: reds for concave and blues for convex regions. (C) Time-dependent rates of actin dynamics (D) Total number of barbed ends depicted as mean (solid curves) and standard deviation (shaded area) over 10 instances of the model. Red curves correspond to the model with LTP induced changes from panel A and green curves to a model without (E) Same for spine volume. Grey markers signal a significant difference between the two conditions (Welch’s t-test with *p <* 0.05 and Bonferroni correction for testing 20 possible time-points per bin)

To model force from actin filaments, we assume that a spine contains a small number of polymerization foci[26], each of them being a branched network of actin-filaments with a certain number of barbed ends *B*. The polymerization at these barbed ends leads to a retrograde flow of the connected filaments through the spine, which, in turn, generates a counter-force – for example by friction or breaking transiently formed bonds. Thus, each focus generates an expansive (outward directed) force, proportional to its number of barbed ends.

To determine the number of barbed ends, we employ a Markov chain in which the current state of a polymerization focus is described by the number of barbed ends *B* and the number of uncapped (exposed) pointed ends *P* (Fig. 1A, left). The actions of different actin binding proteins are then assumed to occur at certain rates, and change the state of the focus accordingly. In particular we consider branching (*B* → *B +* 1) related to the ARP2/3 complex, capping (*B* → *B* − 1) related to the capping protein, severing and depolymerization (*P* → *P* − 1, *B* → *B* − 1) as well as splitting (*P* → *P +* 1, *B* → *B +* 1) both related to the activity of cofilin (see [10, 27] for a review).The number of barbed ends heavily fluctuates and ultimately reaches zero, leading to the removal of the respective focus. Accordingly, also new foci are nucleated randomly such that both the number of active foci and the number of barbed ends at each focus are stochastic. For quantitative analyses of our model, we therefore run multiple simulations and provide average and standard deviations.

To model LTP, we use the relative concentration changes of the respective proteins in Bosch et al. 14 and multiply them with the basal rates in our model (Fig. 1C).

Furthermore, new foci are nucleated randomly with a rate that is also transiently increased during LTP, due to the enhanced availability of actin monomers and severing of cofilin [15].

### Fast decay of LTP-induced perturbations in actin and spine geometry

We then simulated this model for 3.5 hours and tracked the time evolution of spine volume, number of foci and the sum of the number of barbed ends from all foci. In the beginning, the spine evolves from its initial condition (Fig. 1B, *t =* 0 s) into its stationary state given by the membrane force (Fig. 1B, *t =* 1800 s). We observe the typical fast fluctuations of the spine volume, and geometry during the period before the LTP stimulus is applied at *t =* 1h (compare Fig. 1B, *t =* 1800 s and *t =* 3180 s). Upon the onset of the stimulation, we observe a rapid increase in the total number of actively polymerizing barbed ends in the spine (Fig. 1D). However, this increased number of barbed ends decays at the same timescale as the altered rates of the ABPs (dominantly the nucleation rate) and returns to its basal level after around 15 minutes. The volume of the spine (Fig. 1F) undergoes approximately the same time evolution, although somewhat more spread in time, as the spine membrane deforms more slowly. Given that the model can reproduce large volume fluctuation on a timescale of seconds [22], it is not surprising that it cannot preserve information about plasticity events on a longer timescale than a few minutes. Accordingly, significant differences in the volume of a model, with and without LTP-induced modulations of the ABP rates, are only observed for around 30 minutes after the stimulation (compare Fig. 1B, *t =* 5400 s).

When fitting the volume decay (after *t =* 4200*s*) with an exponential function, we obtain a decay-timescale of 523.8 ± 0.2s, which again highlights that our model cannot reproduce altered spine geometries at the timescale of synaptic tagging or early LTP.

This is likely because our model only accounts for a dynamic pool of actin, which is located towards the spine tip and treadmilling very fast and not for the slower treadmilling actin pool – most likely cross-linked actin filaments – that experiments by Honkura et al. had shown. They also already showed that cross-linked filaments are involved in the maintenance of LTP [13]. Yet, they did not quantify whether and how much the stable pool changes after LTP.

### Is there an increase in the stable pool after LTP?

To determine whether the LTP influences the size of the stable pool of actin, we proceeded to wet-lab experiments, focusing on hippocampal cultured neurons, a common model for neuroscience investigations. We used mature neurons (14 days *in vitro*) that expressed a fluorescently-tagged actin variant, carrying a GFP molecule, whose participation in functional reactions has been validated in the past (for example [28], and references therein). We investigated the mobility of the actin molecules using fluorescence recovery after photobleaching (FRAP), a method in which the fluorescent proteins are bleached using a laser beam, in a specific area, and the entry of non-bleached molecules, from neighboring sites, is monitored. Such molecules replace the bleached ones, if they leave the respective area, and the fluorescence signal recovers, with a speed proportional to the molecule mobility [29]. However, if a stable population exists in the respective location, it will not be replaced by mobile molecules, since it does not leave the respective site (in spite of losing fluorescence by bleaching). Such a population is typically termed an immobile fraction, and would correspond, in our conditions, to a stably polymerized actin pool[13].

We applied the FRAP procedure to postsynaptic spines, and determined the recovery of the fluorescence signal over approximately 5 minutes of imaging (Fig. 2A). We then performed the same experiment in cultures subjected to a chemical LTP procedure[30]. We observed a substantially higher immobile fraction at 30 minutes after LTP induction (approx. 45% vs. 20% in the control cultures, Fig. 2B).

**Fig. 2.**
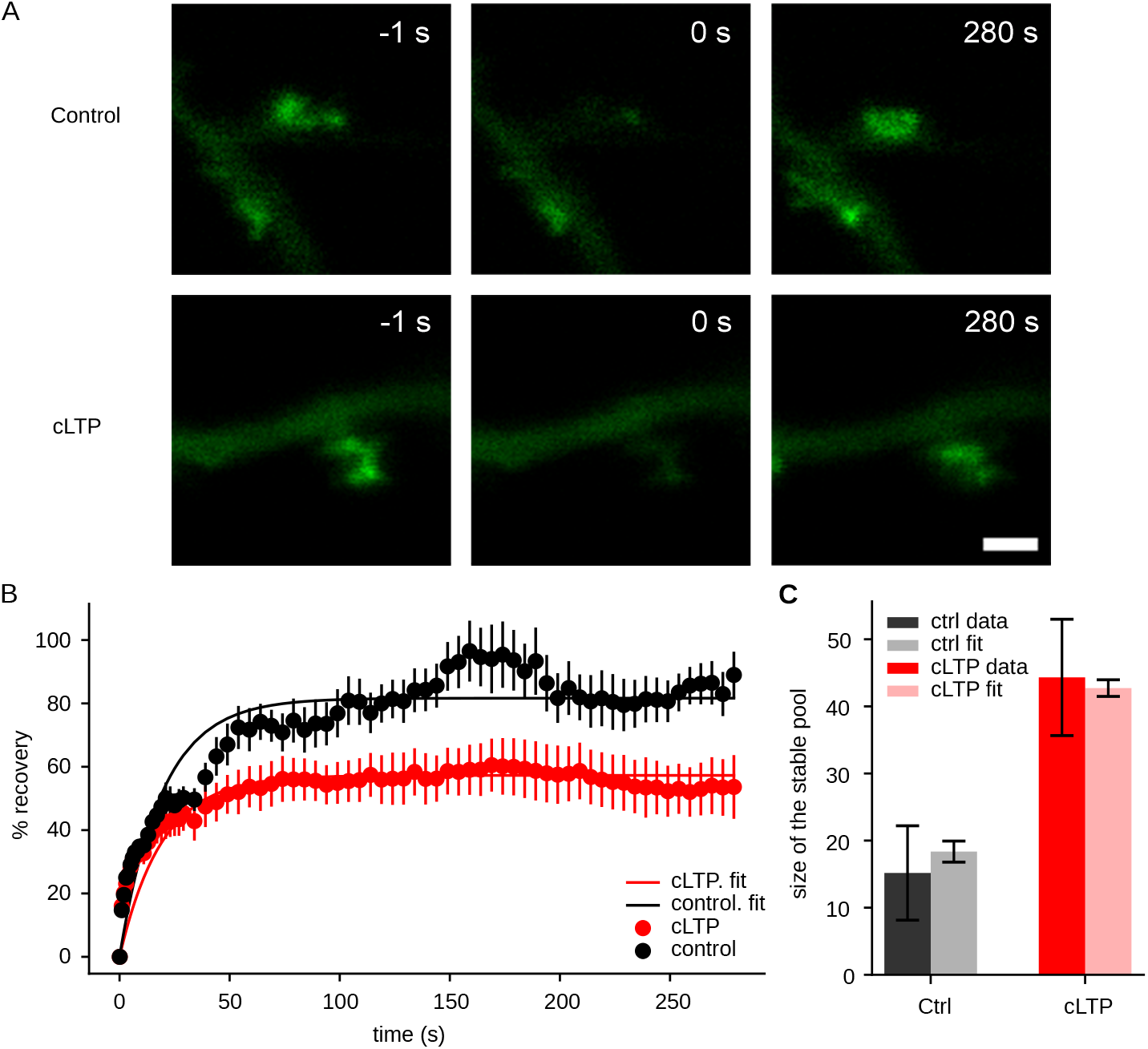
FRAP analysis of actin mobility in dendritic spines. (A) Examples of FRAP experiments performed either without (top panels, control) or 30 minutes after cLTP induction (bottom panels, cLTP). The images show representative dendritic spines before fluorescence photobleaching (left panels, − 1 seconds), immediately afterwards (middle panels, 0 seconds) or after a recovery period (right panels, 280 seconds). (B) An analysis of the fluorescence recovery kinetics, indicated as means ± SEM (*N* = 9 and 10 FRAP experiments for cLTP and control, respectively, from two independent neuronal cultures). The difference in the immobile fractions is significant (*p* = 0.0279, Mann-Whitney ranksum test). (C) Bar graph of the the stable pool fractions in control and 30 minutes after cLTP as measured by mean ± SEM of the immobile fractions in the individual FRAP measurements (solid) and from a curve fit of the curves in panel B (transparent).

We conclude that there is a significant increase in the size of the stable pool, even 30 minutes after LTP induction. Hence, modeling should account for the stable pool and its dynamic, to account for the stabilization of LTP.

### Modeling a stable actin pool

Our next step was therefore to add a time-variant stable pool to our model. This stable pool is assumed to be built-up by cross-linking with new filaments from the dynamic pool at rate *k*_*bind*_ *a*nd decays by unbinding filaments with a rate *k*_*unbind*_. As a measure for the size of the dynamic pool, we use the total number of active barbed ends *B*_*tot*_, such that the dynamic of the stable pool *S* is given by

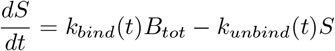

Note that the binding and unbinding rates are time-dependent as cross-linkers detach during the early phase of LTP and reattach after one to five minutes [14, 31]. For simplicity, we model this by a simple step-function (Fig. 3A). Outside of this time window, the stable pool implements a low-pass filtered version of the dynamic actin.

**Fig. 3.**
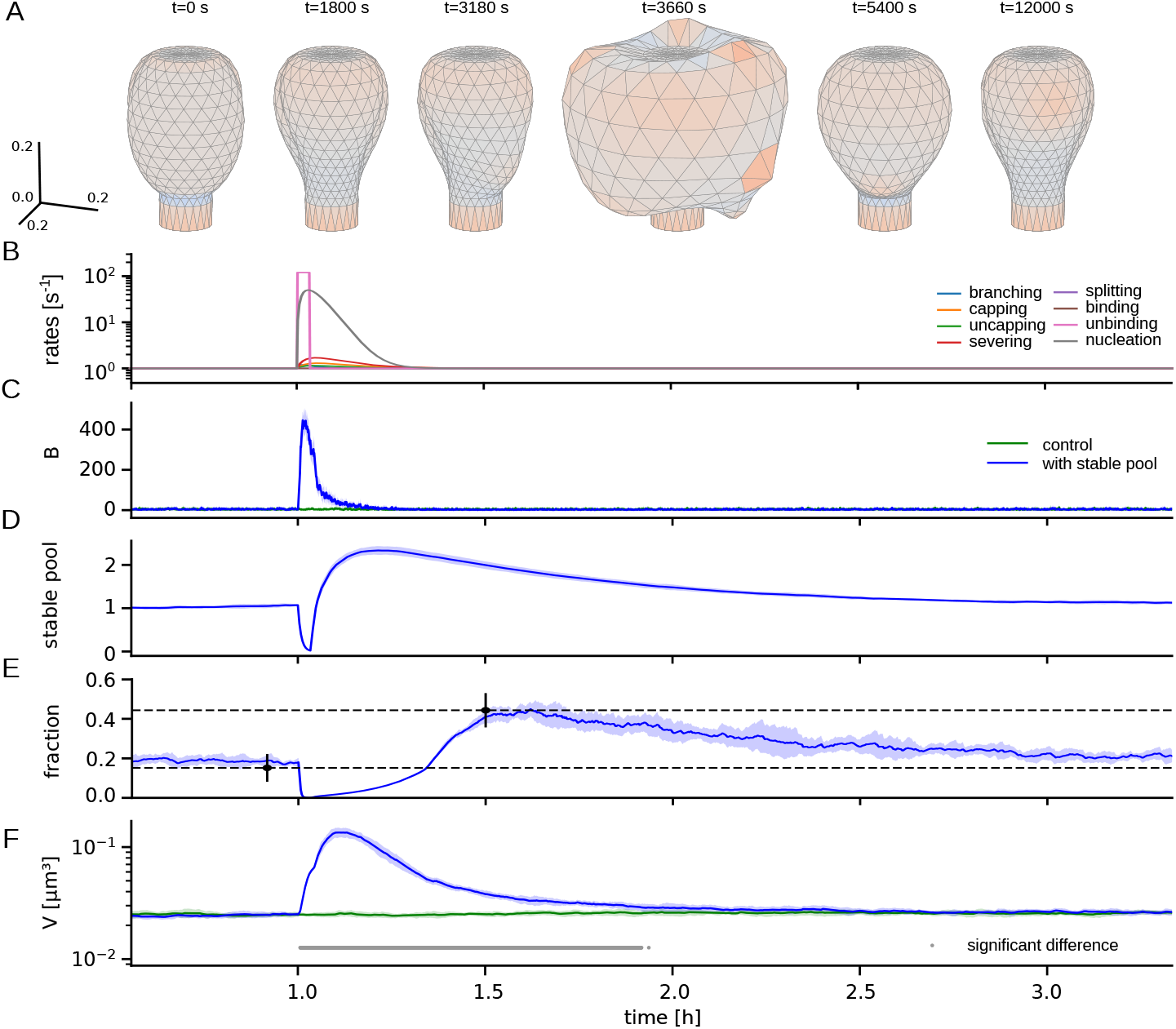
LTP-induced changes in actin dynamics and spine volume persist with novel stable pool. (A) Example spine shapes at different time-points as indicated on top. At *t* = 5400s, that is 30 min after the stimulus, an increased size is still visible. (B) Time-course of rates determining actin dynamics. (C) Total number of barbed ends in the spine (blue). Time average (D) Volume of the spine (blue) and stable pool (green) (E) Fraction of actin allocated to stable pool.

The stable pool also influences the actin force, as a larger stable pool in the spine leads to more resistance against the retrograde movement of the dynamic actin filaments. We implement this by multiplying the dynamic-pool-dependent actin force with *α*_*0*_ (1 + *q* [*S* - *S*_*0*_]_+_). Here, *α*_*0*_ adjusts the overall scaling of the force, *q* scales the stable pool dependent contribution, and *S*_*0*_ acts as an offset for rectification([.]_+_). In this way, the dynamic polymerization foci exert more force, when more stable actin is present, such that their joint action can enlarge the spine.

### Reproduction of overshoot in stable actin ratio

With the stable pool integrated into the model, we simulated the actin dynamic and temporal evolution of spine geometry after LTP. Again, we observe that the time dependent reaction rates after a stimulus (Fig. 3B) induce a large growth in the total number of barbed ends (Fig. 3C). The stable pool drops to very low values once the cross-linkers unbind, but recovers quickly once the reattachment commences (Fig. 3D). Strikingly, as the dynamic actin is still above basal levels at this time, the stable pool overshoots and reaches a level which is higher than its basal state. In the following, this overshoot decays at the slow timescale of the stable pool such that it is visible for 1 to 2 hours. We also evaluated the experimentally measured fraction between stable and total filamentous actin (i.e. stable and dynamic). To calculate the dynamic actin, we use a moving average with a 300 s window on the number of barbed ends, which prevents the fraction to undergo large fluctuations. We find that also the fraction of stable actin exhibits an overshoot, although much later than the stable pool itself (Fig. 3E). This is because the dynamic pool starts higher but decreases faster than the stable pool. Strikingly, the overshoot of the stable pool fraction is highest around the measured time-point, 30 minutes after LTP-stimulation, and matches the experimental measurements (Fig. 3E).

The consequence of the overshoot in stable pool is a longer-lasting significant increase in spine volume (Fig. 3F, grey markers). Hereby, the decay of the LTP-induced increase in spine volume does not seem to follow an exponential decay with a single timescale, but rather has a slow and a fast component. We thus fit the volume as a sum of two exponential decays and obtained a fast timescale of 414.6 ± 0.2 s that accounts for a 350% increase in volume, and a slow one with 2094 ± 5s that accounts for a 50% increase in volume.

We therefore conclude, that the stable actin pool could provide a transient memory of a plasticity event that effects spine geometry for one to two hours – that is, on the timescale of the synaptic tag.

### Model predictions for longevity of synaptic tag

To draw more general conclusions, we also investigate the influence of our model parameters especially on the evolution of the newly introduced stable pool. For this, we consider the situation after the onset of the cross-linker binding and assume that the stable pool starts at zero and the dynamic actin exponentially decays towards its basal value (Fig. 4A, top). Under these assumptions, we can analytically derive the time-course of the fraction of stable actin (Fig. 4B).

**Fig. 4.**
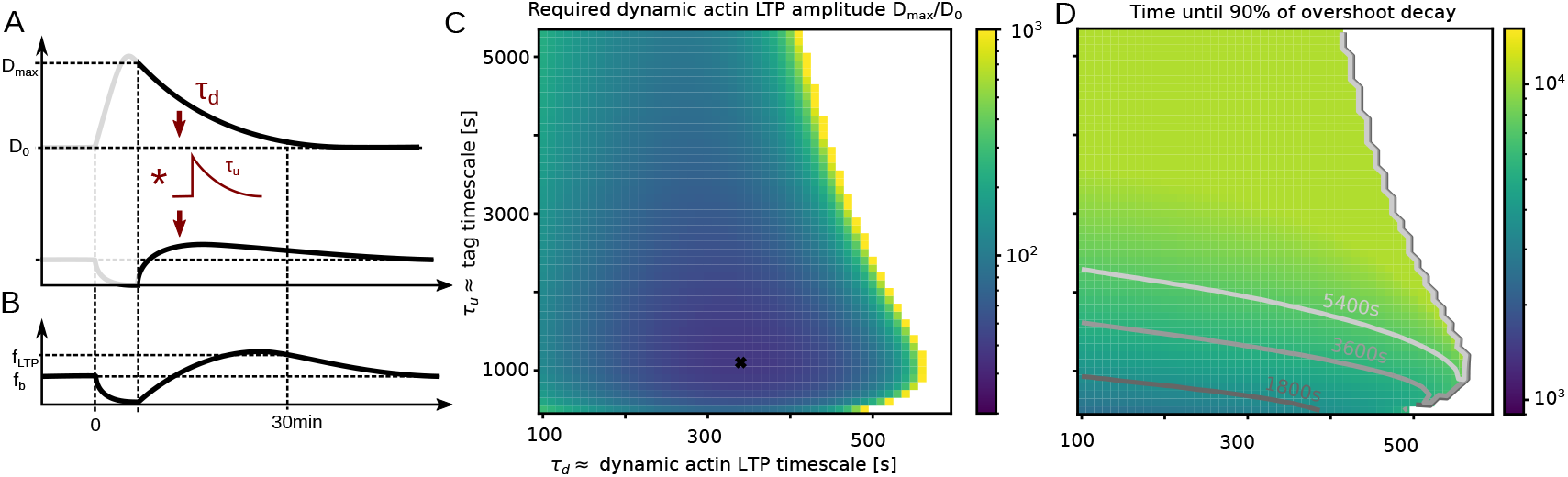
Theoretical predictions. (A) Schematics of theoretical assumptions: When cross-linkers start to bind again, the perturbation in the dynamical pool decays exponentially with *τ*_*d*_, while the stable pool follows it as a leaky integrator with timescale *τ*_*u*_; (B) Prediction for the necessary amplitude in dynamic actin activity (as multiple of the basal dynamic actin activity) measured at the onset of crosslinker binding 2 min after LTP. A minimum at *τ*_*d*_ = 340*s* and *τ*_*u*_ = 1100*s* with *D*_*max*_*/D*_0_ ≈ 38 is observed.

We then systematically vary the cross-linker unbinding timescale *τ*_*u*_ *=* 1/*k*_*unbind*_ and the decay timescale of dynamic actin *τ*_*d*_ and obtain a amplitude of dynamic pool perturbations at the onset of cross-linker binding that is necessary to match the fraction of stable actin from our experiments (see Method section Theoretical Analysis for details). First, we find that matching the data is possible for a broad range of *τ*_*u*_ and *τ*_*d*_ combinations, indicating that the overshoot phenomenon is robust to parameter-variations. However, towards the boundary of possible timescale combinations, the required amplitude of the dynamic pool diverges, which renders these combinations biologically unrealistic. Secondly, we find that the required amplitude has a minimum at *τ*_*d*_ *=* 340*s* and *τ*_*u*_ *=* 1100*s*, where only 40 times more dynamic actin filaments are needed. In all other cases, a larger overshoot before or after 30 minutes is required, which in turn requires more dynamic actin.

Assuming that the dynamic pool has the required amplitude at the onset of cross-linker binding, we then calculated how long the overshoot of the stable pool persists – that is, when its difference to the basal value has decayed to 10% of the maximal value. We find that the longevity of stable pool overshoot is tightly related to the cross-linker unbinding timescale *τ*_*u*_. For sufficiently large *τ*_*u*_, the typical decay timescale of the synaptic tag, of around one to two hours, can be explained by the overshoot of the stable pool.

## Discussion

Using a combined approach of experiments, simulations and theory, we have demonstrated that the stable pool of actin is enlarged 30 minutes after an LTP-like stimulus. Our measurement-constrained model predicts that this enlargement can persist for one to two hours, which is in line with a long-lasting up-regulation of the F-actin content of the spine observed in FRET-experiments [32].

We argue that the stable actin pool could therefore serve as a biophysical implementation of the synaptic tag, as it fulfills all criteria expected for the synaptic tag [5, 7]:

First, a tag needs to be specific for the potentiated synapse, which certainly is the case for the stable pool, as the cross-linked actin filaments stay localized within the spine, as indicated by the FRAP experiments (Fig. 2.

Secondly, the tag should not require protein synthesis, which is also the case for the stable pool, which relies on an increased polymerization of the existing actin molecules. Along this line, actin is one of the most stable proteins in the synapse (half-live of 17-19 days in the mouse brain, more than twice longer than the median value for all proteins;[33]) and there is, to our knowledge, no evidence that actin is synthesized as a plasticity-related protein. Hence, even if protein synthesis is already active in our experiments 30 minutes after LTP, we can assume that the actin dynamics would be unaltered.

Third, the tag should decay within one to two hours, which we demonstrated here. Nonetheless, our theoretical analysis further shows that the experimental measurements can be reproduced by a variety of timescales in the same regime as the ones used here. Hence, further experiments will be needed to draw more quantitative conclusions.

Fourth, the tag should be able to recruit newly-synthesized proteins. As stable actin affects the geometry of the spine, it will also regulate the space available to reorganize the PSD. Moreover, the PSD itself interacts with actin [34, 35] and consequently also with the stable pool. However, our model does not yet cover a mechanism by which protein capture is realized and gives rise to functional plasticity at the synapse, but this is definitely one of the next steps to take.

Apart from its role for a synaptic tag, the inclusion of a stable actin pool is a novelty in comparison to previous models [19, 21–23], which have largely neglected the time-evolution of the stable pool. Yet, as also these models rely on barbed end numbers or densities, the modifications presented here for the stable pool could be similarly applied to them. Functionally, our stable pool acts as a slow (low-pass filtered) version of the fast dynamic actin. Synapses with such a fast-and-slow dynamics have been shown to exhibit many properties of brain-like learning and memory [36] and may help with continual learning [37]. Accordingly, the synapse model we introduced can be assumed to exhibit similar beneficial characteristics for learning and memory.

In summary, our work therefore strongly suggests that interaction between spine geometry and time-dependent actin polymerization and cross-linking plays an essential role in the consolidation of synaptic plasticity and memory, by providing a long-lasting, but transient memory of past stimuli, which is consistent with a biophyiscal implementation of the synaptic tag.

## Acknowledgments

This work was funded by the German Science Foundation under CRC1286 “Quantitative Synaptology”, projects C03 and A03. We would like to thank Simon Dannenberg, Stefan Klumpp, Jannik Luboeinski, Francesco Negri, Christian Tetzlaff and Florentin Wörgötter for fruitful discussions on the project.

## Author contributions

Performed Experiments: CB, SR; Model Conception: MT, MF; Simulations: MT; Acquired Funding MF, SR; Conceptualization MF, SR; Wrote Manuscript: MT, CR, SR, MF

## (Online) Methods

### Experimental model and study participant details

We used newborn rats (*Rattus norvegicus*) to prepare cultures. Animals were handled according to the regulations of the local authorities, the University of Göttingen and the State of Lower Saxony (Landesamt für Verbraucherschutz, LAVES, Braunschweig, Germany). All animal experiments were approved by the local authority, the Lower Saxony State Office for Consumer Protection and Food Safety (Niedersächsisches Landesamt für Verbraucherschutz und Lebensmittelsicherheit) and were performed in accordance with the European Communities Council Directive (2010/63/EU).

### Experimental procedures: wet lab

#### Rat hippocampal cultures

We prepared dissociated cultures from hippocampi following procedures established before, exactly as described in [28]. In brief, hippocampi (from newborn wild-type, Wistar animals) were dissected in HBSS (Hank’s Buffered Salt Solution, 140mM NaCl, 5mM KCl, 4mM, 6mM glucose, NaHCO_3_, 0.4mM KH_2_PO_4_ and 0.3mM Na_2_HPO_4_). They were then incubated with an enzyme solution, prepared in DMEM (Dulbecco’s Modified Eagle Medium, #D5671, Sigma-Aldrich, Germany), for 60 minutes. The enzyme cocktail contained 50mM EDTA, 100mM CaCl_2_,0.5 mg/mL cysteine and 2.5 U/mL papain, and was carbogen-saturated for 10 min before application. The materials were then exposed to an enzyme-deactivation buffer (DMEM with 0.2 mg/mL bovine serum albumin, 0.2 mg/mL trypsin inhibitor and 5 percent FCS, fetal calf serum). After cellular trituration, we seeded the material on 18mm circular coverslips, at 80,000 cells per coverslip. The coverslips were prepared by treating with nitric acid, followed by sterilization and exposure to 1 mg/mL poly-L-lysine, overnight. The cells adhered to the coverslips for 1–4 h at 37^°^C DMEM containing horse serum (10 percent), 2mM glutamine and 3.3mM glucose. This medium was then replaced with Neurobasal-A (Life Technologies, Carlsbad, CA, USA) containing 2% B27 (Gibco, Thermo Fisher Scientific, USA) supplement, 1% GlutaMax (Gibco, Thermo Fisher Scientific, USA) and 0.2% penicillin/streptomycin mixture (Biozym Scientific, Germany). The coverslips were incubated at 37^°^C, under 5% CO_2_ atmosphere, until use.

DIV 4 primary hippocampal neurons were transfected in order to express GFP-tagged actin, using Lipofectamine^™^ 2000 Transfection Reagent (Invitrogen^™^) as vehicle.

#### Chemical LTP induction

The procedure of inducing chemical LTP was adapted from the methods described by Zheng and collaborators [30]. Briefly, DIV15 neurons were washed once with a basal buffer (150mM NaCl, 5mM KCl, 2mM CaCl_2_, 2mM MgCl_2_, 10mM HEPES, 30mM D-Glucose, pH=7.34-7.36) to get rid of the debris from the cell media. Then, neurons were washed twice with Mg^2+^-free buffer (150mM NaCl, 5mM KCl, 2mM CaCl_2_, 10mM HEPES, 30mM D-Glucose, 0.02mM bicuculline and 0.001mM picrotoxin, pH=7.34-7.36), for a total of 5 minutes. Neurons were further incubated at 37^°^C in a glycine-supplemented buffer (150mM NaCl, 5mM KCl, 2mM CaCl_2_, 10mM HEPES, 30mM D-Glucose, 0.02mM bicuculline, 0.001mM picrotoxin and 0.2 mM glycine, pH=7.34-7.36) for 15 minutes. Following a wash with the basal buffer, the neurons were incubated at 37°C in Mg^2+^ buffer (150mM NaCl, 5mM KCl, 2mM CaCl_2_, 2mM MgCl_2_, 10mM HEPES, 30mM D-Glucose, 0.02mM bicuculline and 0.001mM picrotoxin, pH=7.34-7.36) for 30 minutes, prior to live imaging in basal buffer.

#### Neuronal live imaging

Individual neuronal spines were used for the FRAP (Fluorescence Recovery After Photobleaching) experiments. A TCS SP5 confocal microscope (Leica, Wetzlar, Germany) equipped with an HCX Plan Apochromat 100×1.40 oil immersion objective was used for the imaging. The 488 nm wavelength of an Argon laser was used for imaging of GFP.. Before bleaching, 4 images were taken, and then, the region of interest was bleached for 100 ms. The bleaching intensity was defined as in [28]: 50*µ*W, at 488 nm. After bleaching, 10 images were taken every 1 s, then 10 images every 2 s, and 50 images every 5 s. For the control condition, DIV15 hippocampal neurons were shortly washed with the basal buffer and directly live-imaged, as described above.

#### FRAP image analysis

The FRAP movies were analyzed using a custom-written routine in MATLAB (The Math-Works Inc, Natick, MA, USA). The images were loaded, and were then automatically aligned, and the FRAP area was determined automatically, by comparing the last frame before bleaching to the first one post-bleaching, and was set as the FRAP region of interest (ROI). The ROI intensity was then determined for all of the frames, was corrected for background, by subtracting the average intensity signals in other, non-cellular areas, and was then normalized to the pre-bleaching intensity. The resulting FRAP curves were then averaged and displayed as mean and SEM in Fig 2.

FRAP experiments have been conducted for *N =* 9 spines (each from an independent FRAP experiment) from two cultures for the control and *N =* 10 spines (again, each from an independent FRAP experiment) from two cultures for the cLTP case. As described, values were normalized by the basal fluorescence of each spine and then depicted by their mean and standard error of the mean.

The experimentally-derived immobile (stable) fractions represent the % of the initial intensity of the spines that did not recover by the end of the recording. This definition fits well with the observation that the recovery curves saturated within our observation time.

The fractions of the stable pool were also analyzed by fitting an exponential decay

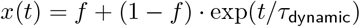

to both curves simultaneously using pythons `curve fit` function. Here, *f =* (*f*_*b*_, *f*_*LT P*_)^*T*^ are the fractions of stable actin and *τ*_*dynamic*_ the timescale of the dynamic pool. The standard deviation of the fit fractions was obtained from the covariance matrix

### Biophyiscal model of membrane dynamics

#### Membrane model and initialization

The spine membrane is modeled with a 3D triangular mesh. It is initialized as a sphere with a radius of 0.25*µ*m and 20 segments along the latitudinal and longitudinal direction. Every ring along the latitudinal axis is alternatingly rotated by ± 9^°^, to obtain regular triangles. We then flatten the top four rings of vertices to the level of the fourth ring and declare these points as the PSD. Further, to model the spine neck, we replace the bottom 2 rings by a cylinder with the same radius as the second-last ring, and a length of 0.1. Afterwards, the *x* and *y* coordinates of all points are scaled by a factor of 0.7 to obtain an initial configuration corresponding to a stubby spine.

#### Membrane dynamic

The vertices of the PSD and the neck remain fixed, as these regions are supported by either transmembrane proteins like neurexins/neuroligins (PSD) or by a rigid ring-structure of actin filaments (neck). At all other locations, a vertex *i’*s position 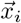 changes according to the forces generated by the actin filaments 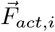 and the forces generated by the membrane 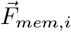 at that vertex. For numerical stability, we added a force 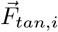 which moves vertices along the spine surface to preserve shape but obtain an approximately equilateral mesh. The movement of vertex *i* is then given by:

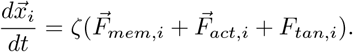

The following sections detail the calculations of all three forces.

#### Membrane force

Following previous works ([21–23], see [38] for a review), we derive the membrane force from the Canham-Helfrich free energy

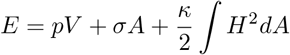

where *p* is the pressure difference between inside and outside the membrane, V is the spine (head) volume, *σ* the elasticity of the membrane, *A* its area, *κ* the membrane bending modulus and *H* the mean curvature. The force is the derivative of this energy with respect to the (three dimensional) position of the mesh vertices 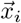:

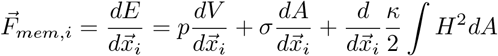

In the following we will derive approximations for the three terms on the right hand side for our mesh:

*volume* The volume can be calculated as

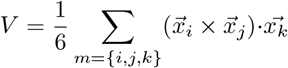

where the sum runs over all triangles *m* with points *i, j* and *k*. For the contribution of the individual triangle *V*_*m*_, the derivatives are

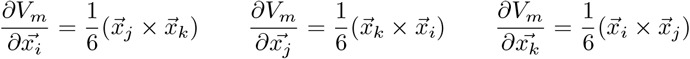

In the source code, we calculate these three contributions for each triangle and then add them to the summed force of the respective vertices *i, j* and *k*.

*area* The area can be calculated as

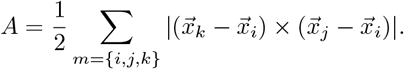

Consider triangle sites 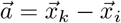 and 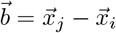, then

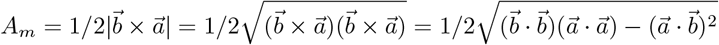

through the Binet-Cauchy-identity.

We know that 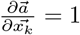 and 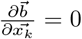, such that we can calculate the derivative as

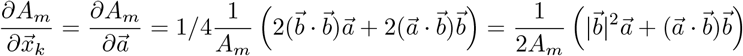

In our implementation, we calculate the area contributions by iterating through all triangles. Due to symmetry reasons, we do not calculate the derivatives w.r.t 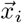 and 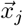 but just calculate the above contribution for each cyclic permutation of *i, j* and *k* and add this contribution to the area-dependent force of the respective vertex.

*curvature* We approximate the curvature integral (different from [21], but see [22]) as

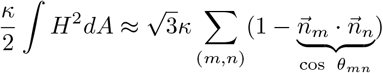

with *m* and *n* being adjacent surfaces of the mesh and 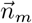 and 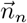 their (unit length) face normals. These normals, in turn, depend on the position of the vertices spanning the triangle.

In the following, we assume that the points *i, j* and *k* span a triangle *m* and *l* is the third point in the neighboring triangle *n* containing *i* and *j* such that we can use site 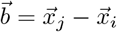 twice. The (unit length) face normals are then

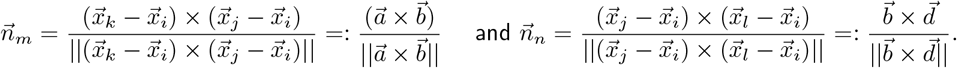

We apply Binet-Cauchy to the numerator 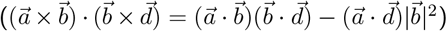 and to the denominator 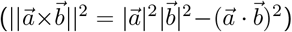 obtaining a product of denominators

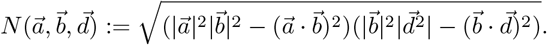

With this, we can easily evaluate the derivatives:

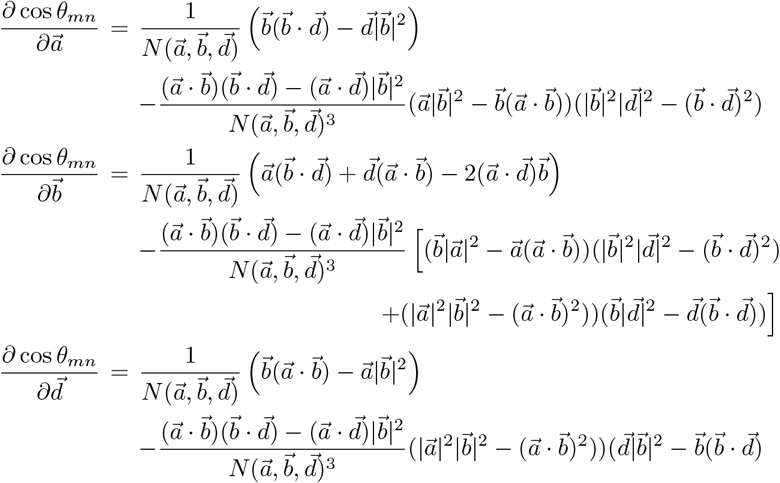

Then, with 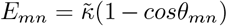 and 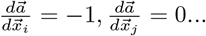 the derivatives w.r.t. each vertex are:

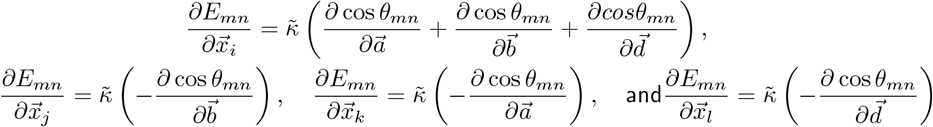

In the code, the vectors 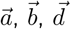 as well as the scalars 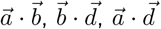 and 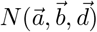 are pre-calculated for all triangles before the force contributions are calculated and assigned to the respective vertices.

These terms signify the force contribution of triangles *m* and *n* (that share the edge *ij*) to the vertices *i, j, k, l*. By evaluating them for all edges and summing the force contributions from different triangle constellations for each vertex, we obtain the total curvature-induced force for each vertex.

#### Tangential force and mesh equilibration

The above described numerical approximations work best if all involved triangle edges are approximately equal. Therefore, we induce an additional tangential force. For this, we first sum the vectors of all edges emerging from a vertex *i* into a sum-vector *S*_*i*_. Thus, if the edges are not equal, the sum vector will point towards the long edges, where the point should be moved. We then determine to which of triangles containing point *i* the sum-vector can be projected (positive scalar products with both triangle sites starting at *i*) and project it to that plane. We finally multiply the sum vector by a factor of 10*pN/µm* to obtain a tangential force 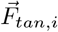 Note, this tangential force moves the vertices only along the existing surface but still can slightly change volume, area and curvature, especially for strongly curved local neighborhoods.

### Actin dynamics and forces

Actin forces are generated from distinct polymerization foci within the spine [21, 26]. Hence our model allows for multiple foci of actin activity which act on the membrane at different points.

### Dynamic of one focus

We abstract the complex growth of the filament tree to a Markov process with two state variables: the number of barbed ends *B* and the number of uncapped (exposed) pointed ends *P*. During the simulation, there are multiple transitions which can take place corresponding to the actions of various actin binding proteins:

▪ *Branching* (*B* → *B +* 1): Due to the attachment of ARP2/3 a new filament can branch off an existing one which gives rise to a new barbed end. The rate at which this happens is

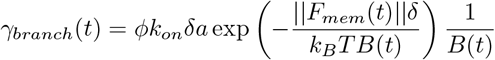
▪ where *ϕ* is a proportionality constant, *k*_*on*_ is the G-actin assembly rate constant, *a* is the available actin concentration, *F*_*mem*_ is the counteracting membrane force, *δ* is the length of G-actin, *k*_*B*_*T* is the thermal energy, and *B*(*t*) the number of barbed ends at time t. (one quantity time dependent)
▪ *Capping* (*B* → *B* − *1*): When a capping protein (e.g. CapZ) attaches to a barbed end, it stops further polymerization, essentially removing it.
▪ *Severing* (*B* → *B* − *1, P* → *P* − *1*): Cofilin binding leads severing of a filament followed by its complete depolymerization.
▪ *Splitting* (*B* → *B +* 1, *P* → *P +* 1): Cofilin binding also under some conditions, leads to the further growth of the severed filament to an active filament.
▪ *Uncapping* (*P* → *P +*1): The site where ARP2/3 is bound is also a capped pointed end. When it unbinds, an uncapped pointed end is formed, which is free to then depolymerize.

To indicate the status variables of different foci, we will use a index in the following, that is *B*^(*z*)^ is the number of barbed ends from focus *z*. Note, once the number of barbed ends reaches 0, we assume that there is no more filament to branch and the focus is removed from the simulation. Thus, it is also necessary that new foci are nucleated.

#### Nucleation

New actin foci within the spine which are formed according to a nucleation rate *γ*_*nucl*_. The location for this – the nucleation points – are chosen probabilistically: First, a set of *n* points is generated which are at 80% of the distance between the membrane points and the origin. Second, the distance *d*_*j*_ of each of these points to the center of the PSD is calculated. Third, one of these point is selected as nucleation point with probability 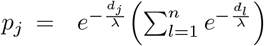, where *λ* is a scaling parameter for the PSD distance. The growth direction of the actin focus 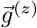 is calculated as a unit vector along the line between origin and the nucleation point. If the membrane moves such that the nucleation point is outside the membrane, we move it backward along this direction vector until it is inside the spine again.

#### Actin force

We assume that each focus has a spatial extent orthogonal to its growth direction described by a Gaussian kernel with amplitude *α* and spread *σ* as

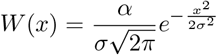

To calculate the force contribution of focus *z*, we first select all vertices 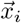 that lie in the growth direction from the focus origin. We then consider a line originating from the focus origin and extending in the growth direction 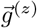 and calculate the orthogonal distance of these points 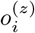.

The actin force from focus *z* onto vertex *i* is then determined by this orthogonal distance and the barbed ends:

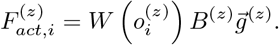

The contributions of each focus are summed at all vertices.

### Modeling LTP by time-dependent rates

The concentration of ABPs vary in a time-dependent manner in response to an LTP stimulus [14]. This time development can be fit with a double exponential function defined by two time constants - a rise time *τ*_*1*_ and a fall time *τ*_*2*_:

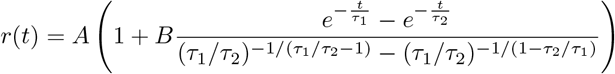

Here, *A* is the basal rate and *B* is the percentage by which the rate changes. The values of these have been obtained from fitting measurements in [14] and are reported in Table 1

**Table 1.** Time-dependent model parameters varied through LTP

| Process/protein | A | B | $\tau_1(s)$ | $\tau_2(s)$ | Comment/Source |
| --- | --- | --- | --- | --- | --- |
| Branching/ARP2/3 | 2.0 | 0.2 | 120 | 180 | [21] and [14] |
| Capping/CapZ | 1 | 0.3 | 120 | 360 | [21] and [14] |
| Uncapping/ARP2/3 | 1/30 | 0.15 | 100 | 300 | [21] and [14] |
| Severing/Cofilin | 1 | 0.7 | 120 | 300 | [21] and [14] |
| Splitting/Cofilin | 1 | 0.7 | 120 | 300 | [14] |
| Nucleation | 0.05 | 69 | 20 | 240 | adapted to experiment |

### Model of the stable pool

We assume that filaments from dynamic pool transit to the stable pool through cross-linking with rate *k*_*b*_. Cross-linker unbinding leads to a shrinkage of the stable pool, and is proportional to its size and an unbinding rate *k*_*u*_. Thus we have

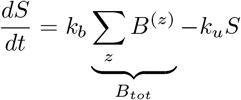

Therefore, our stable pool model behaves as an exponential low-pass filter of the total number of barbed ends with a timescale *k*_*u*_. Upon LTP-inducing stimulations, the cross-linker unbinding rate is instantly changed to *k*_*u*_ *=* 120 times the basal value and reset after 2 minutes.

The stable pool moreover also influences spine geometry. Assuming that it hinders retrograde movement, the expansive force from the dynamic actin filaments should proportionally grow with its size. We therefore replace the constant *α* in the kernel for our force equation by

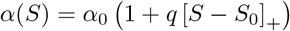

where *α*_*0*_ scales the stable pool independent force contributions and *q* the stable pool influence. [.]_+_ marks a rectifications such that *S*_*0*_ sets a minimum amount of cross-linked actin at which an influence on the expansive forces can be observed.

### Model parameters

We listed all time-constant parameters in Table 2. In comparison to [21, 22], we decreased the spine size to obtain realistic volumes and adapted a few parameters along that line.

**Table 2.**
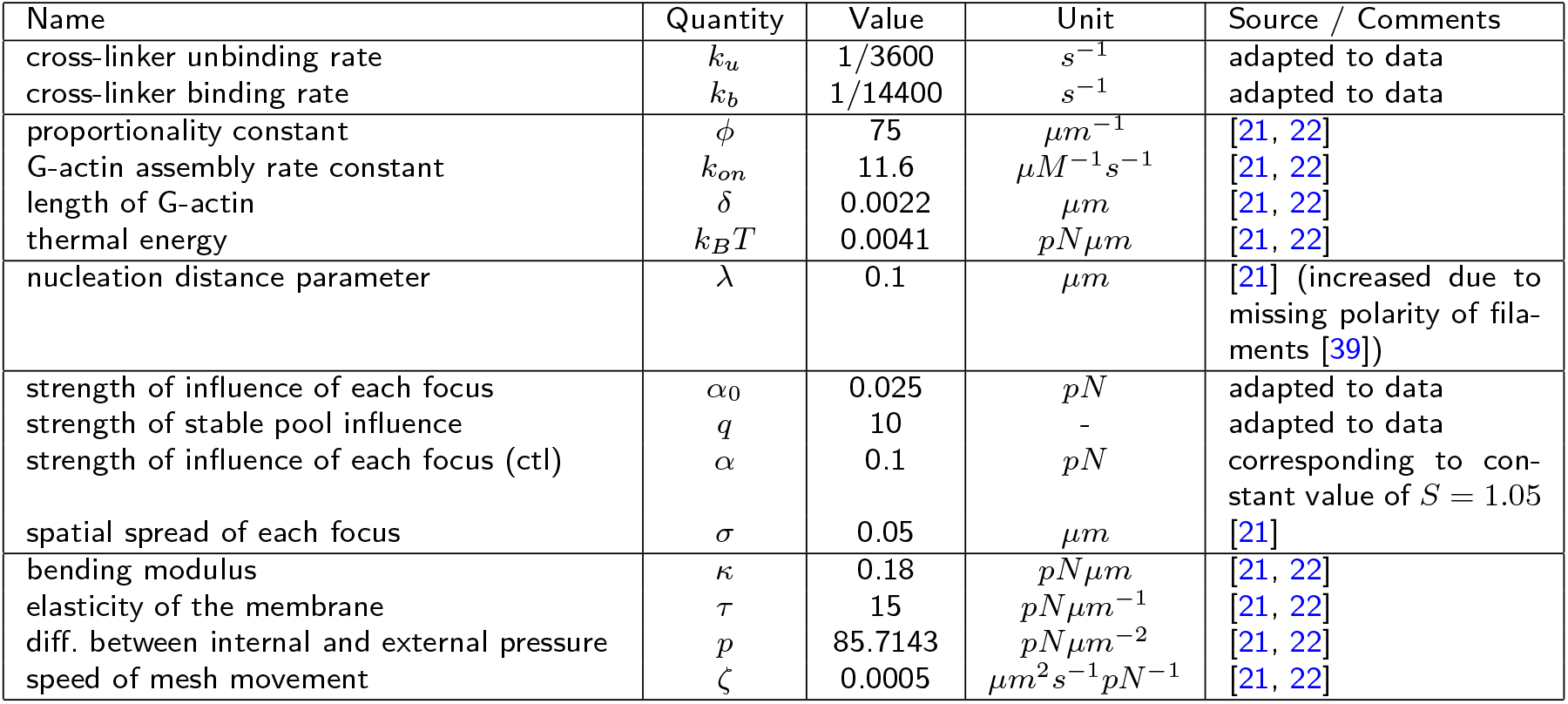
Time-independent model parameters

#### Simulations and statistxical analysis

Simulations have been conducted and analyzed in `python` and `numpy`. For each condition (control, without and with stable pool), we conduct *n =* 10 simulation and depict mean and standard deviation values. To determine whether the mean is significantly increased for the LTP cases as compared to the control condition, we obtain the statistics for a (one-sided) Welch-t-test for each time point, combine them into groups of 20 time-points and see whether one of them exhibits a p-value of *p <* 0.05/20 (significance level for Bonferroni correction for multiple testing)

### Theoretical Analysis

In the following we will derive an analytical solution for the time-course of the stable pool and the necessary conditions for dynamic actin for matching the experimentally measured actin fraction.

We assume that, after crosslinkers start binding again, the dynamic pool *D* (which is proportional to the number of barbed ends) follows a time-course:

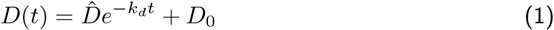

while the stable pool follows

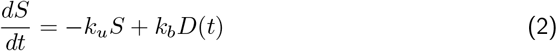

and is assumed to start at a value of *S*(0) = 0. This is an initial value problem based on a first order inhomogeneous linear ODE. The solution to the homogeneous equation (*D*(*t*) = 0) is

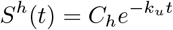

By variation of the constant *C*, we obtain a solution to the inhomogeneous case:

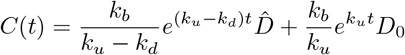

and arrive at

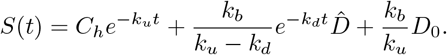

Using the initial value *S*(0) = 0, we obtain

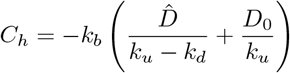

Hence, the resulting time-course is a difference of exponentials.

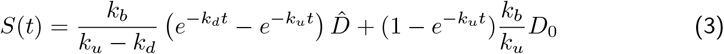

We next want focus on the stable pool fraction of the overall actin

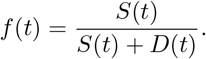

For the basal value at *t* → ∞, this equates to

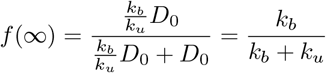

Thus, for a given (or measured) basal value of *f*(∞) = *f*_*b*_, we can constrain the ratio between binding and unbinding rate of the cross-linkers:

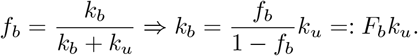

Using this relation in the time-course results in

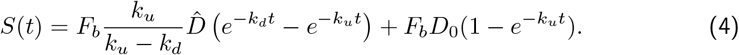

Secondly, we measured the fraction after LTP *f*(*t*_*LT P*_ *=* 30min) = *f*_*LT P*_, which results in

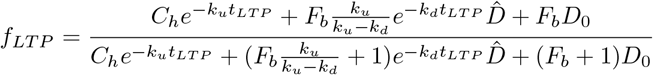

or

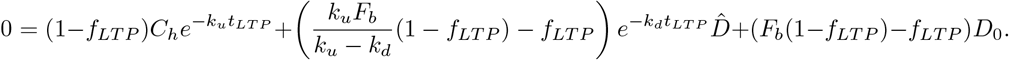

Substituting 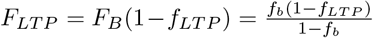 and converting to unit-less *D*_*0*_ = 1,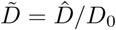:

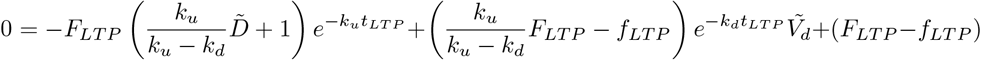

These equations can give us 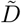 for given *k*_*u*_ and *k*_*d*_, which tells us how much larger the dynamic actin must be compared to basal state after cross-linkers bind again:

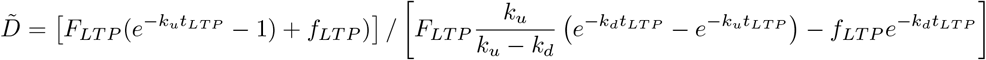

Finally, we use this value 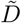 in equation 4 and numerically calculate the resulting timecourse of the stable pool and the stable pool fraction. From this time-course we determine the maximum value and the time-point at which the difference to the basal value has decayed to 10% of that of the maximum value, which we depict as the persistence time of the synaptic tag in Fig 4C.

## References

[1] Bliss, T.V., Lømo, T.: Long-lasting potentiation of synaptic transmission in the dentate area of the anaesthetized rabbit following stimulation of the perforant path. The Journal of physiology 232(2), 331–356 (1973)

[2] Bliss, T.V., Collingridge, G.L.: A synaptic model of memory: long-term potentiation in the hippocampus. Nature 361(6407), 31–39 (1993)

[3] Abraham, W.C., Jones, O.D., Glanzman, D.L.: Is plasticity of synapses the mechanism of long-term memory storage? NPJ science of learning 4(1), 9 (2019)

[4] Dringenberg, H.C.: The history of long-term potentiation as a memory mechanism: Controversies, confirmation, and some lessons to remember. Hippocampus 30(9), 987–1012 (2020)

[5] Frey, U., Morris, R.G.: Synaptic tagging and long-term potentiation. Nature 385(6616), 533–536 (1997)

[6] Abraham, W.C.: How long will long-term potentiation last? Philosophical Transactions of the Royal Society of London. Series B: Biological Sciences 358(1432), 735–744 (2003)

[7] Redondo, R.L., Morris, R.G.: Making memories last: the synaptic tagging and capture hypothesis. Nature Reviews Neuroscience 12(1), 17–30 (2011)

[8] Martin, K.C., Kosik, K.S.: Synaptic tagging—who’s it? Nature Reviews Neuroscience 3(10), 813–820 (2002)

[9] Krucker, T., Siggins, G.R., Halpain, S.: Dynamic actin filaments are required for stable long-term potentiation (LTP) in area CA1 of the hippocampus. Proceedings of the National Academy of Sciences 97(12), 6856–6861 (2000)

[10] Pollard, T.D.: Actin and actin-binding proteins. Cold Spring Harbor perspectives in biology 8(8), 018226 (2016)

[11] Shaw, J.E., Kilander, M.B., Lin, Y.-C., Koleske, A.J.: Abl2: cortactin interactions regulate dendritic spine stability via control of a stable filamentous actin pool. Journal of Neuroscience 41(14), 3068–3081 (2021)

[12] Okamoto, K., Bosch, M., Hayashi, Y.: The roles of CaMKII and F-actin in the structural plasticity of dendritic spines: a potential molecular identity of a synaptic tag? Physiology 24(6), 357–366 (2009)

[13] Honkura, N., Matsuzaki, M., Noguchi, J., Ellis-Davies, G.C., Kasai, H.: The subspine organization of actin fibers regulates the structure and plasticity of dendritic spines. Neuron 57(5), 719–729 (2008)

[14] Bosch, M., Castro, J., Saneyoshi, T., Matsuno, H., Sur, M., Hayashi, Y.: Structural and molecular remodeling of dendritic spine substructures during long-term potentiation. Neuron 82(2), 444–459 (2014)

[15] Borovac, J., Bosch, M., Okamoto, K.: Regulation of actin dynamics during structural plasticity of dendritic spines: Signaling messengers and actin-binding proteins. Molecular and Cellular Neuroscience 91, 122–130 (2018)

[16] Fonseca, R.: Activity-dependent actin dynamics are required for the maintenance of long-term plasticity and for synaptic capture. European Journal of Neuroscience 35(2), 195–206 (2012)

[17] Pinho, J., Marcut, C., Fonseca, R.: Actin remodeling, the synaptic tag and the maintenance of synaptic plasticity. IUBMB life 72(4), 577–589 (2020)

[18] Matsuzaki, M., Honkura, N., Ellis-Davies, G.C., Kasai, H.: Structural basis of longterm potentiation in single dendritic spines. Nature 429(6993), 761–766 (2004)

[19] Miermans, C., Kusters, R., Hoogenraad, C., Storm, C.: Biophysical model of the role of actin remodeling on dendritic spine morphology. PloS One 12(2), 0170113 (2017)

[20] Alimohamadi, H., Bell, M.K., Halpain, S., Rangamani, P.: Mechanical principles governing the shapes of dendritic spines. Frontiers in Physiology 12, 657074 (2021)

[21] Bonilla-Quintana, M., Wörgötter, F., Tetzlaff, C., Fauth, M.: Modeling the shape of synaptic spines by their actin dynamics. Frontiers in synaptic neuroscience 12, 9 (2020)

[22] Bonilla-Quintana, M., Wörgötter, F., D’Este, E., Tetzlaff, C., Fauth, M.: Reproducing asymmetrical spine shape fluctuations in a model of actin dynamics predicts self-organized criticality. Scientific reports 11(1), 4012 (2021)

[23] Bonilla-Quintana, M., Rangamani, P.: Biophysical modeling of actin-mediated structural plasticity reveals mechanical adaptation in dendritic spines. Eneuro 11(3) (2024)

[24] Bonilla-Quintana, M., Wörgötter, F.: Exploring new roles for actin upon LTP induction in dendritic spines. Scientific reports 11(1), 7072 (2021)

[25] Fukazawa, Y., Saitoh, Y., Ozawa, F., Ohta, Y., Mizuno, K., Inokuchi, K.: Hippocampal LTP is accompanied by enhanced F-actin content within the dendritic spine that is essential for late LTP maintenance in vivo. Neuron 38(3), 447–460 (2003)

[26] Frost, N.A., Shroff, H., Kong, H., Betzig, E., Blanpied, T.A.: Single-molecule discrimination of discrete perisynaptic and distributed sites of actin filament assembly within dendritic spines. Neuron 67(1), 86–99 (2010)

[27] Van Troys, M., Huyck, L., Leyman, S., Dhaese, S., Vandekerkhove, J., Ampe, C.: 23 Ins and outs of adf/cofilin activity and regulation. European journal of cell biology 87(8-9), 649–667 (2008)

[28] Reshetniak, S., Ußling, J.-E., Perego, E., Rammner, B., Schikorski, T., Fornasiero, E.F., Truckenbrodt, S., Köster, S., Rizzoli, S.O.: A comparative analysis of the mobility of 45 proteins in the synaptic bouton. The EMBO Journal 39(16), 104596 (2020)

[29] Axelrod, D., Koppel, D., Schlessinger, J., Elson, E., Webb, W.W.: Mobility measurement by analysis of fluorescence photobleaching recovery kinetics. Biophysical journal 16(9), 1055–1069 (1976)

[30] Zheng, N., Jeyifous, O., Munro, C., Montgomery, J.M., Green, W.N.: Synaptic activity regulates ampa receptor trafficking through different recycling pathways. Elife 4, 06878 (2015)

[31] Sekino, Y., Tanaka, S., Hanamura, K., Yamazaki, H., Sasagawa, Y., Xue, Y., Hayashi, K., Shirao, T.: Activation of n-methyl-d-aspartate receptor induces a shift of drebrin distribution: disappearance from dendritic spines and appearance in dendritic shafts. Molecular and Cellular Neuroscience 31(3), 493–504 (2006)

[32] Okamoto, K.-I., Nagai, T., Miyawaki, A., Hayashi, Y.: Rapid and persistent modulation of actin dynamics regulates postsynaptic reorganization underlying bidirectional plasticity. Nature neuroscience 7(10), 1104–1112 (2004)

[33] Fornasiero, E.F., Mandad, S., Wildhagen, H., Alevra, M., Rammner, B., Keihani, S., Opazo, F., Urban, I., Ischebeck, T., Sakib, M.S., et al.: Precisely measured protein lifetimes in the mouse brain reveal differences across tissues and subcellular fractions. Nature communications 9(1), 4230 (2018)

[34] Kuriu, T., Inoue, A., Bito, H., Sobue, K., Okabe, S.: Differential control of postsynaptic density scaffolds via actin-dependent and-independent mechanisms. Journal of Neuroscience 26(29), 7693–7706 (2006)

[35] Chen, X., Jia, B., Zhu, S., Zhang, M.: Phase separation-mediated actin bundling by the postsynaptic density condensates. Elife 12, 84446 (2023)

[36] Benna, M.K., Fusi, S.: Computational principles of synaptic memory consolidation. Nature neuroscience 19(12), 1697–1706 (2016)

[37] Kaplanis, C., Shanahan, M., Clopath, C.: Continual reinforcement learning with complex synapses. In: International Conference on Machine Learning, pp. 2497–2506 (2018). PMLR

[38] Guckenberger, A., Gekle, S.: Theory and algorithms to compute helfrich bending forces: a review. Journal of Physics: Condensed Matter 29(20), 203001 (2017) 10.1088/1361-648X/aa6313

[39] Tatavarty, V., Kim, E.-J., Rodionov, V., Yu, J.: Investigating sub-spine actin dynamics in rat hippocampal neurons with super-resolution optical imaging. PloS one 4(11), 7724 (2009)

